# Singing training predicts increased insula connectivity with speech and respiratory sensorimotor areas at rest

**DOI:** 10.1101/793083

**Authors:** A.M. Zamorano, R.J. Zatorre, P. Vuust, A. Friberg, N. Birbaumer, B. Kleber

## Abstract

The insula contributes to the detection of salient events during goal-directed behavior and participates in the coordination of motor, multisensory, and cognitive systems. Recent task-fMRI studies with trained singers suggest that singing experience can enhance the access to these resources. However, the long-term effects of vocal training on insula based networks are still unknown. In this study, we employed resting-state fMRI to assess experience-dependent differences in insula co-activation patterns between conservatory-trained singers and non-singers. Results indicate enhanced bilateral insula connectivity in singers relative to non-singers with constituents of the speech sensorimotor network in both hemispheres. Specifically, with the cerebellum (lobule VI, crus 2), primary somatosensory cortex, the parietal lobes, and the thalamus. Furthermore, singing training predicted enhanced bilateral co-activation of primary sensorimotor areas representing the larynx (with left dorsal anterior insula, dAI) and the diaphragm (with bilateral dAI)—crucial regions for cortico-motor control of complex vocalizations, as well as the thalamus (with bilateral posterior insula/left dAI) and the left putamen (with left dAI). Together, these data support a crucial role of the insula in respiratory and laryngeal control and suggest that singing experience enhances the integration of somatosensory information within the speech motor system, perhaps by strengthening salient associations of bodily signals associated with conscious and non-conscious aspects of expressive language production within a musical framework.

## INTRODUCTION

Research over the last two decades has provided compelling evidence that the extensive practice and performance of fine motor, sensory, and multimodal integration skills required for mastering a musical instrument can have profound effects on the functional and structural organization of the brain (Klein et al., 2016; Schlaug, 2015; Vuust et al., 2014; Zatorre, 2013). As a widely-used framework for studying human neuroplasticity, musical training has shown that adaptive changes in the brain are linked to the rehearsed musical action, encompass quantifiable behavioral correlates (i.e., improved performance), occur in response to short and long-term training, and appear to be more pronounced when formal training began during a sensitive period before the age of seven years (Herholz and Zatorre, 2012; Munte et al., 2002). Professionally trained singers, on the other hand, have rarely been studied in this context. They differ from instrumentalists as they (i) may commence with formal training significantly later (Donahue et al., 2014; Schirmer-Mokwa et al., 2015) and (ii) optimize a motor system for music production that has already undergone substantial maturation throughout speech motor development and informal singing experience (Smith, 2006; Stadler Elmer, 2011; Weiss-Croft and Baldeweg, 2015). As professional singers still require a large amount of deliberate practice to meet the musically prescribed standards, the current study used conservatory trained singers as a model for testing experience-dependent neuroplasticity within the speech-motor system (Kleber and Zarate, 2014; Zarate, 2013).

The neural pathways underlying basic vocalizations have been thoroughly described in primates (Jurgens, 2002, 2009), whereas higher-level aspects of vocal motor production have been studied in the context of human speech (Price, 2012). Together, they revealed a hierarchically organized structure-function relationship in which basic vocal patterns generated in the brain stem and the periaqueductal gray interact with limbic regions, and the higher-level control of complex volitional vocalizations is performed in the cortex (Hage and Nieder, 2016; Simonyan and Horwitz, 2011). The interactions between these regions in the context of speech motor processes are described and formalized by neurocomputational models (Guenther, 2016; Parrell et al., 2019). One prominent model (DIVA, Directions Into Velocities of Articulators) suggests that the dorsal speech-motor stream engages a sensory integration interface consisting of auditory, ventral somatosensory, and inferior parietal cortices, which in turn interacts via the arcuate fasciculus with the articulatory motor network located in ventral primary- and pre-motor regions (Guenther and Hickok, 2015). In addition, the supplementary motor area, the basal ganglia, and the cerebellum take part via cortico-thalamic loops and contribute to feedback and feedforward transformations that enable the accurate performance of planned vocalizations (Dick et al., 2014; Hertrich et al., 2016).

The insular cortex, on the other hand, is not prominently featured in these models, although it has long been considered to contribute to motor aspects of speech production (Ardila et al., 2014; Dronkers, 1996; Price, 2000) due to its involvement in vocalization-related autonomic and perceptual-motor processes (Craig, 2002, 2009) and structural connectivity with constituents of the dorsal sensorimotor stream (Hickok, 2017; Remedios et al., 2009; Riecker et al., 2008). Indeed, the insular cortex connects to a vast number of cortical and subcortical brain regions subserving sensory, emotional, motivational and cognitive processes (Gogolla, 2017; Nomi et al., 2018). Taking a centrally located position within these networks, its broad functional roles involve the conscious and non-conscious integration of bodily information (Craig, 2002, 2009), sympathico-vagal (homeostatic) regulation of emotions (Strigo and Craig, 2016), the detection and assessment of salient sensory inputs (Berret et al., 2019; Singer et al., 2009), as well as the coordination of large-scale network dynamics underlying higher-level cognitive processes (Menon and Uddin, 2010; Seeley et al., 2007; Uddin, 2015). With respect to speech, clinical and functional imaging suggest a bilateral insula contribution to linguistic processes and a more left-lateralized contribution of the dorsal anterior insula to non-linguistic aspects of speech production and perception (Ackermann and Riecker, 2010; Oh et al., 2014). Singing-related perceptual and motor processes, in contrast, have indicated a right-hemispheric dominance (Ackermann and Riecker, 2004; Jeffries et al., 2003; Riecker et al., 2000), perhaps due to the processing of prosodic or melodic elements of vocalizations (Oh et al., 2014), similar to reports of brain asymmetries for speech and melody perception in the auditory cortex (Albouy et al., 2020). This notion is supported by neuroimaging studies with trained classical singers, in which the right anterior insula seems to contribute to the stabilization of vocal pitch in the presence of sensory feedback perturbations (Kleber et al., 2017; Kleber et al., 2013). However, the long-term effects of vocal training on insula-based networks remain unknown.

The current study employed resting-state (rs-)fMRI to quantify experience-dependent differences in large-scale insular connectivity between trained singers and non-singers, following the notion that rs-fMRI reflects the record of repeated task-dependent synchronized activation between brain regions (Guerra-Carrillo et al., 2014). To this end, a seed-based approach was used (Fox and Raichle, 2007) to explore temporally correlated spontaneous low-frequency blood oxygenation level-dependent fluctuations between the whole brain and a-priori defined regions of interests (ROI). Following a tripartite insula subdivision model (Deen et al., 2011), ROIs encompassed the left and right posterior insula (PI), dorsal anterior insula (dAI), and ventral anterior insula (vAI) (Nomi et al., 2018; Uddin et al., 2014). We predicted enhanced insula connectivity in singers compared to non-singers, specifically between the AI with regions of the dorsal sensorimotor speech stream involved in vocal tract coordination (Kleber et al., 2017; Kleber et al., 2013), analogous to prior findings with musicians trained in playing an instrument (Zamorano et al., 2017). A high level of task-relevant structure function specificity would provide further evidence for a central role of the insula in vocal motor control and may help describing the effect of vocal experience on insula-based sensory integration networks.

## METHODS

### Participants

A total of 25 right-handed subjects without reported history of neurological or psychiatric disease participated in this study. Participants were subdivided in two groups based on their singing expertise. Professional singers (n= 12, 6 female, 32.7 ± 8.6 yrs) with music conservatory education took their first formal singing lesson at the average age of 16 years (± 6.7) and accumulated an average of 12957 hours of singing experience (range: 1456-38220). Non-singers (n=13, 6 female, 28.1 ± 7.3 yrs) consisted of University of Tübingen medical school students, who neither received prior singing training nor reported any involvement in occasional singing activities (e.g., choirs, informal rock bands etc.). All participants were informed about the details of the study and provided written consent. The study was conducted under a protocol approved by the research ethics board of the University of Tübingen.

### Behavioral experiments

#### Pitch-matching accuracy

Prior to rs-fMRI, a behavioral pitch-matching task was performed to assess experience-dependent differences in singing accuracy between trained singers and non-singers. During this task, a total of 54 pseudorandomized musical target intervals were presented via headphones using Max/MSP software to control the experiment (Cycling 74, San Francisco, California, USA). Upon each presentation, participants were prompted to reproduce the pitch of two tones with their singing voice. The first tone always started at the fundamental frequency of 311.13 Hz for females (D#4 in musical notation) and 155.565 Hz for males (D#3 in musical notation). The second tone was either the same (4x) or differed from the first tone by one (6x), three (12x), five (12x), six (12x) or seven (8x) semitones, with an equal number of ascending and descending intervals. Each tone was played with a duration of 900-ms, separated by a 200-ms gap. Vocal reproduction was recorded and saved in wave format for offline automated analyses of pitch-matching accuracy.

Pitch-matching accuracy was defined by the deviation between the target tones presented via headphones and the tones sung by the participants. In a first step, the deviation was estimated in cents (one semitone corresponds to 100 cents) using a custom-made script within the CUEX performance analysis system (Friberg et al., 2005) runing in Matlab (The MathWorks, Inc., Natick, Massachusetts, United States). In a second step, statistical analyses were performed to assess the effect of singing expertise on pitch-matching accuracy. As the 54 pitch-matching responses per subject and the responses for same interval sizes could not be regarded as independent observations, a generalized linear mixed model was fit using the lmer function provided by the lme4 package (Bates et al., 2015) in R (v.3.6.1; R Development Core Team, 2019). The full model included ‘group’ (singers *vs* non-singers) as fixed effect, and ‘subject’ and ‘interval size’ as random effects [m_full <-lmer(pitch_devation ∼ group + (1|subject_nr) + (1|interval_size), REML=FALSE)]. The statistical significance of the full model was assessed from Chi square distribution and p values estimated with a likelihood ratio test (Barr et al., 2013) by comparing the full to a null model without the fixed factor ‘group’ [m_null <-lmer(pitch_devation ∼ 1 + (1|subject_nr) + (1|interval_size), REML=FALSE)]. As the residuals were found to be non-normal and there was evidence of heteroscedasticity of variances, the data were log transformed (log10) before statistical analysis. We confirmed that the transformed values produced comparable results to non-transformed values.

### Resting-state fMRI acquisition

Resting-state data were acquired over a period of 7.5 minutes with the eyes closed. Participants were instructed to stay awake and to think of nothing in particular. Magnetic resonance imaging was performed using a 3-Tesla whole body MRI Scanner (Siemens MAGNETOM Prisma™ 3T, Erlangen, Germany). For each subject, 225 echo-planar volumes were acquired (repetition time, 2000 ms; echo time, 30 milliseconds; matrix dimensions, 64 x 64; field of view, 1260 mm; 30 transversal slices; slice thickness, 4 mm; flip angle, 90 degrees) The structural imaging data for anatomical reference consisted of T1-weighted images (mprage, repetition time, 2300 milliseconds; echo time, 4.18 milliseconds; matrix dimensions, 512 x 512; field of view, 256 x 256 mm; 176 slices; slice thickness, 1 mm; flip angle, 9 degrees).

### fMRI data preprocessing

Following our previous approach (Zamorano et al., 2017; Zamorano et al., 2019), functional images were preprocessed with the Data Processing Assistant for Resting-State fMRI (DPARSF; Chao-Gan and Yu-Feng, 2010), based on the Statistical Parametric Mapping software package (SPM8; http://www.fil.ion.ucl.ac.uk/spm) and the Data Processing & Analysis of Brain Imaging toolbox (DPABI; http://rfmri.org/DPABIDPARSF_V3.1_141101). The first 10 volumes from each data set were discarded prior to preprocessing. Following slice-time correction and co-registration, gray and white matter were segmented from co-registered T1 images using the unified segmentation model (Ashburner and Friston, 2005). The resulting parameter file was used to normalize the functional images (3mm3 voxel size) to standard Montreal Neurological Institute (MNI) stereotactic space, which were subsequently smoothed with an isotropic Gaussian kernel (FWHM: 6mm_3_). Nuisance regression parameters included white matter, CSF, and the six head motion parameters. WM and CSF masks were generated using SPM’s tissue probability maps (empirical thresholds: 90 % for WM mask and 70 % for CSF mask). No global signal regression was performed to avoid introducing BOLD signal distortions (Murphy et al., 2009). Head motion was below 2.0 mm maximum displacement or 2.0° of any angular motion across all participants. A temporal filter (0.006–0.1 Hz) was applied to reduce low frequency drifts and high frequency physiological noise.

### Functional connectivity analysis

Statistical analyses were performed in SPM8 to assess voxel-wise connectivity maps of posterior, ventral anterior and dorsal anterior insula ROIs in each hemisphere. The six insula ROIs consisted of parcellated insular subdivisions (posterior, PI; dorsal anterior, dAI; and ventral anterior insula, vAI) in MNI stereotactic space based on the clustering of functional connectivity patterns during resting state. The ROI template images were kindly provided by Dr. Ben Deen (https://bendeen.com/data/).

The main connectivity patterns across participants (N=25) were first determined by entering z-transformed connectivity maps into one-sample t-tests for each insula ROI (Figure 1). Significance threshold for voxel-wise statistics was set to p<0.05 familywise error corrected (FWER) to validate our results against previously published patterns (Deen et al., 2011).

**Figure 1:**
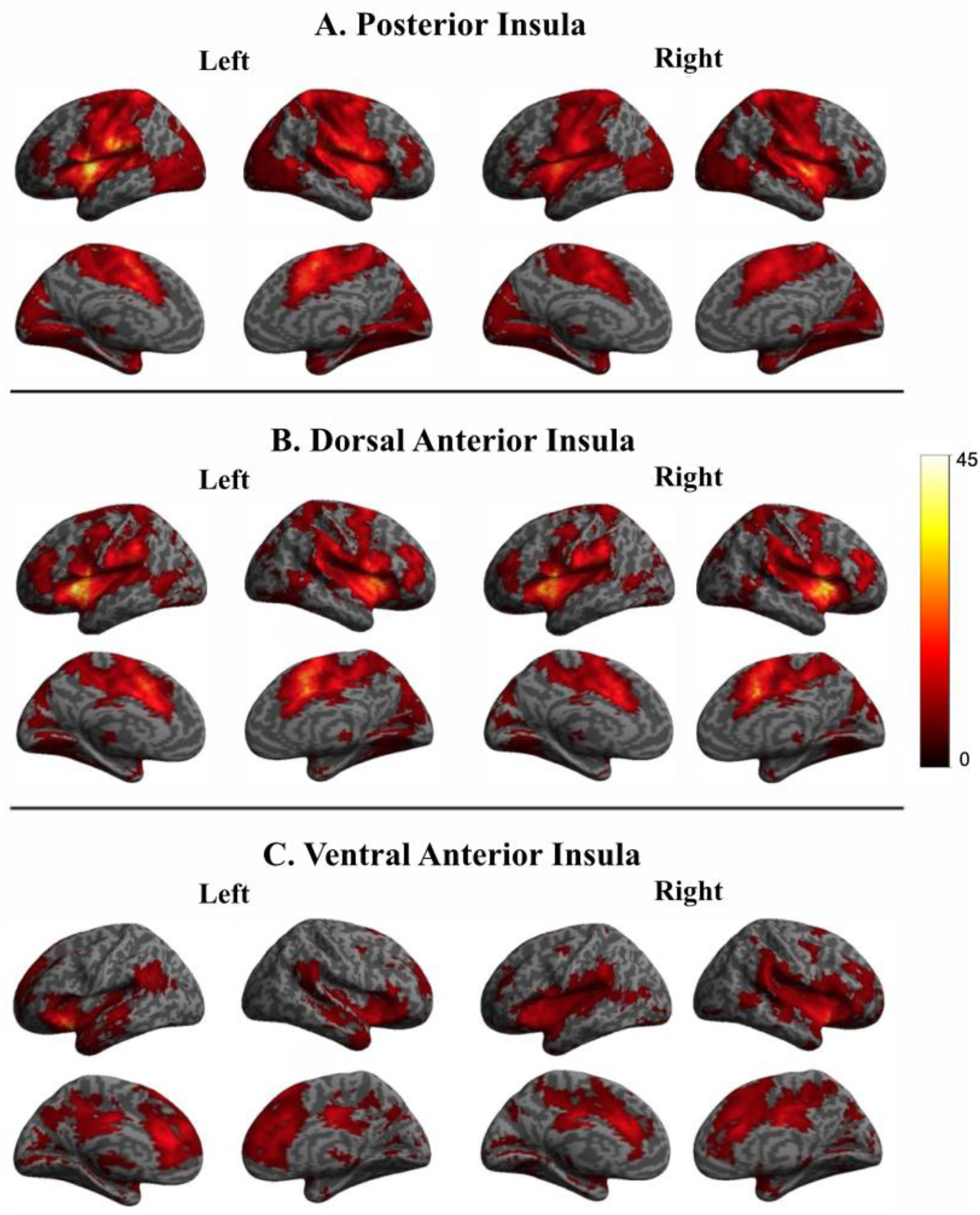
Main patterns of functional connectivity across all participants for ventral anterior insula (vAI), dorsal anterior insula (dAI), and posterior insula (PI) subdivisions, showing both overlapping and distinct co-activated regions, consistent with a more “cognitive” dAI and a more “emotional” vAI pattern, as well as a stronger focus on somatosensory cortical networks for the PI. Significance thresholds were set at P<0.05, familywise error (FWER) corrected. Connectivity maps for left hemisphere seeds are shown on the left; maps for right hemisphere seeds are on the right.

Subsequently, voxel-wise differences between insula-connectivity maps were computed between singers and non-singers using independent two-sample t-tests for each insula ROI. A cluster-extent based thresholding method was employed to increase sensitivity for detecting true activations in studies with moderate sample sizes (Woo et al., 2014). This method accounts for the fact that individual voxel activations are not independent of the activations of their neighboring voxels in spatially smoothed data (Heller et al., 2006) and is hence used to effectively reduce the possibility of obtaining false positive clusters while improving the degree of confidence in inferences about specific voxels (Woo et al., 2014).

Monte Carlo simulation was as implemented using DPABI’s instantiation (Yan et al., 2016) of AlphaSim (Cox, 1996) and the DPABI gray matter mask to determine cluster-extent thresholds at *P* < 0.05 FWER correction for multiple comparisons. Following the recommendations for cluster-extent thresholding in fMRI analyses (Woo et al., 2014), a stringent primary voxel-level threshold of *P*< 0.001 was applied using smoothness estimation based on the spatial correlation across voxels to avoid inaccurate FWER correction. Only clusters surviving the FWER probability threshold were used for statistical inference. T-values of significantly activated peak-voxels within clusters are presented as MNI coordinates. Effect sizes using Cohen’s d (*d* = 2t / sqrt(df)) were computed and subsequently adjusted [unbiased Cohen’s d; *d*_*unbiased*_ = d (1 – (3 / 4df -1))] to control for overestimation of the effect size due to the moderate sample size (Cohen, 2013).

The Automated Anatomical Labeling atlas of Tzourio-Mazoyer (aal, Tzourio-Mazoyer et al., 2002) was used to determine anatomical regions. For regions already cytoarchitectonically mapped, we used the Anatomy Toolbox (Eickhoff et al., 2005).

### Regression analysis

Two regression analyses were performed in SPM8 to correlate insula ROI connectivity maps with (i) pitch-matching accuracy across all participants (deviation from target pitch, log-transformed) and (ii) accumulated singing training in professional singers. Cluster extent FWER correction was applied as detailed above.

## RESULTS

### Pitch-matching accuracy

All individuals were able to complete the pitch matching task without difficulty. The mean deviation from target pitch across pitch-matching trials was 17.5 cent in singers (SD = 9.71, Median = 16) and 72.8 in non-singers (SD = 103, Median = 29.5). A linear mixed effects model (group = fixed effect; subjects and interval size = random effect) using log-transformed pitch-accuracy data (S-Figure 1) indicated that singing expertise significantly improved pitch-matching accuracy (χ2(1)= 5.80, p=0.015). Re-transformation of the data in this model showed that deviation from target pitch was about 21.5 cent ±1.3 (SD) lower in singers (15.2 cent) compared to non-singers (36.7).

### Voxel-wise functional connectivity of insula subdivisions

#### Main connectivity patterns across all participants

Details of whole-brain connectivity patterns for the PI, dAI, and vAI (left and right, respectively) across all participants are shown in Figure 1 and were consistent with those reported earlier (Deen et al., 2011; Uddin et al., 2014).

The left and right *posterior insula* (PI) connectivity pattern (Figure 1A) involved the ipsi- and contralateral insula subdivisions and adjacent operculae but was mostly centered on sensorimotor regions (S1, S2, M1, and pre-motor cortices, SMA and pre-SMA), frontal speech motor areas (IFG, Broca’s area), the posterior and mid cingulate cortex (PCC and MCC), the temporal (superior temporal, Heschl’s, and fusiform gyrus), parietal and (superior and inferior, supramarginal gyrus, and the precuneus) occipital lobe (calcarine, lingual, and cuneus), and the cerebellum.

The left and right *ventral anterior insula* (vAI) connectivity pattern (Figure 1C) involved ipsi- and contralateral insula subdivisions, adjacent rolandic and frontal operculae, as well as bilateral regions in the frontal lobe (DLPFC, inferior frontal gyrus, IFG) and frontal limbic cortices (ventro anterior prefrontal cortex, VAPFC; orbitofrontal gyrus, OFC), the cingulate cortex (ACC and MCC), SMA and pre-SMA, temporal (e.g., Heschl’s gyrus, temporal pole, superior, middle and inferior temporal gyrus) and the parietal lobes (supramarginal and angular gyrus, precuneus).

The left and right *dorsal anterior insula* (dAI) connectivity pattern (Figure 1B) involved other ipsi- and contralateral insula subdivisions, adjacent rolandic and frontal operculae, as well as bilateral sensorimotor regions in the primary (S1) and secondary (S2) somatosensory and motor (M1) cortices, pre-motor cortex and supplementary motor area (SMA), the occipital (calcarine, lingual, and cuneus), temporal (superior temporal gyrus, Heschl’s and fusiform gyrus, temporal pole) and parietal areas (supramarginal, superior and inferior parietal gyrus, and the precuneus), the cerebellum, as well as the inferior frontal (IFC), dorsolateral prefrontal (DLPFC), and cingulate cortex (posterior, PCC; middle, MCC, and anterior, ACC).

Subcortically, all three insula sub-regions showed co-activation patterns with the basal ganglia (putamen, pallidum and caudate) and the thalamus.

### Differences in insula connectivity between trained singers and non-singers

T-contrasts of connectivity maps between singers and non-singers revealed increased functional connectivity for each insula subdivision in singers (Figure 2 and Table 1). Reversed comparisons yielded no significant differences.

**Table 1:**
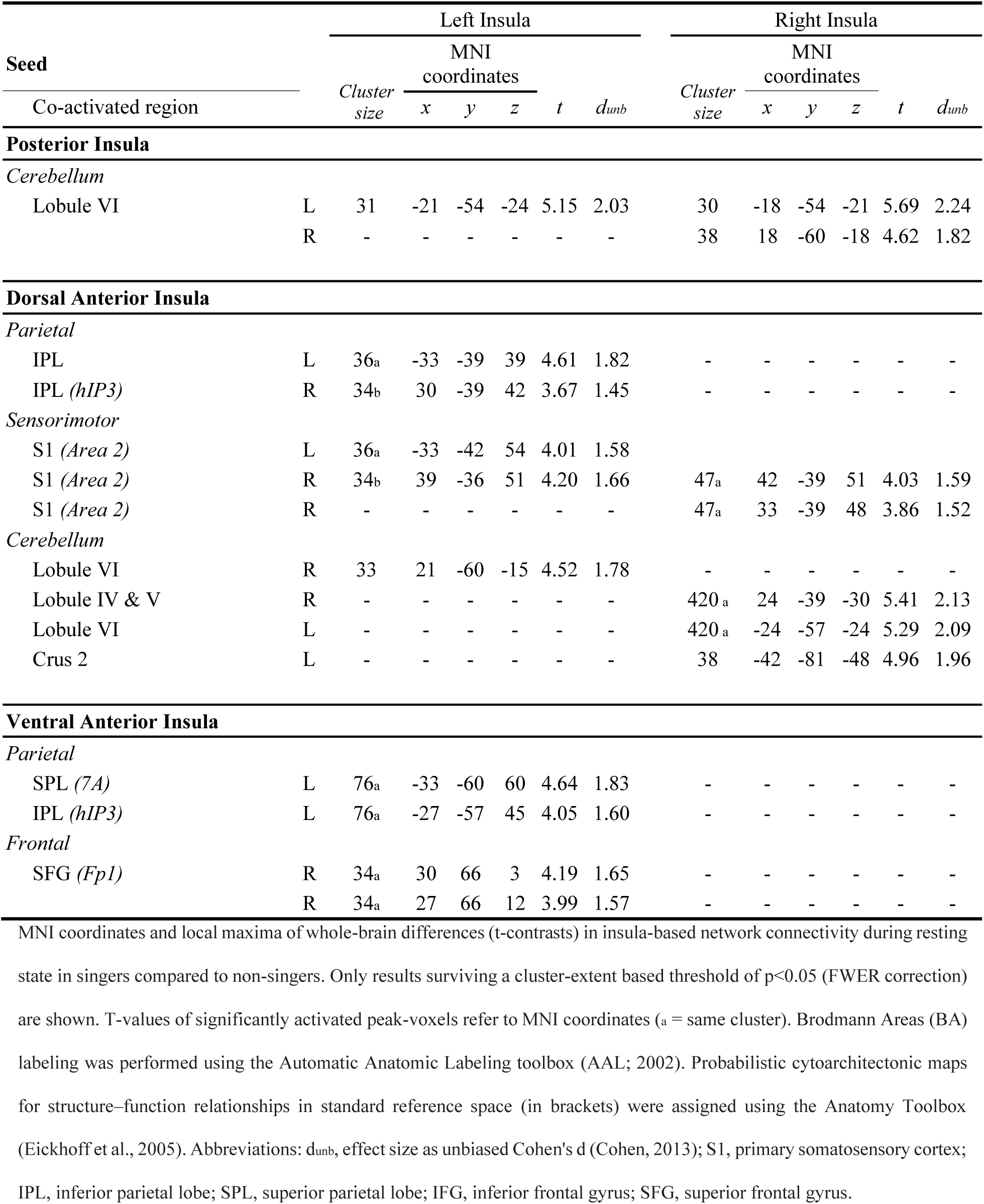
Regions with increased insula connectivity in singers compared to non-singers

**Figure 2:**
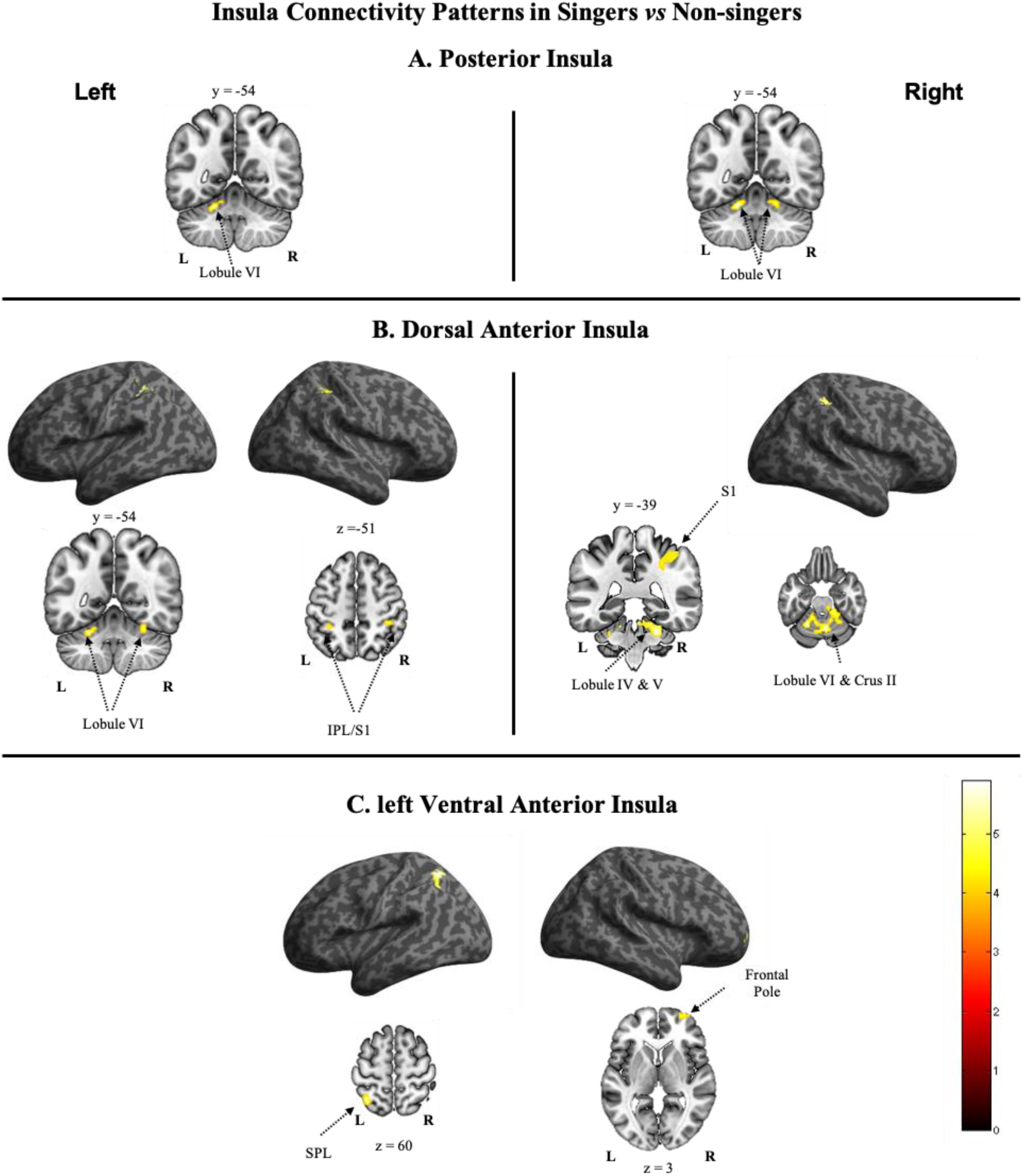
T-maps showing significant group differences in functional insula connectivity during resting-state in singers. Significance thresholds using a cluster-extent based thresholding method for between-group differences were set at p>0.05 (FWER). Detailed information is provided in Table 1. Abbreviations: IPL, inferior parietal lobe; SPL, superior parietal lobe; S1, primary somatosensory cortex; IV-VI and Crus II, cerebellar lobules.

The **left and right PI** (Figure 2A and Table 1) showed increased functional connectivity with lobule VI of the cerebellum in singers.

The **left dAI** (Figure 2B, Table 1) showed increased functional connectivity with bilateral inferior parietal and primary somatosensory cortex, the cerebellum (lobule VI) and the thalamus. The **right dAI** showed increased functional connectivity with the cerebellum (lobule VI and Crus 2) and the right S1. Connectivity with the right inferior frontal gyrus failed to reach significance (Broca’s homologue, Area 45).

The **left vAI** (Figure 2C, Table 1) showed increased functional connectivity with the left superior and inferior parietal cortex. The **right vAI** did not reveal significant differences.

### Regression Results

Multiple regression analyses were performed to determine correlations between insula ROI connectivity maps and behavioral variables: (i) pitch-matching accuracy and (ii) accumulated singing training (Table 2).

**Table 2.**
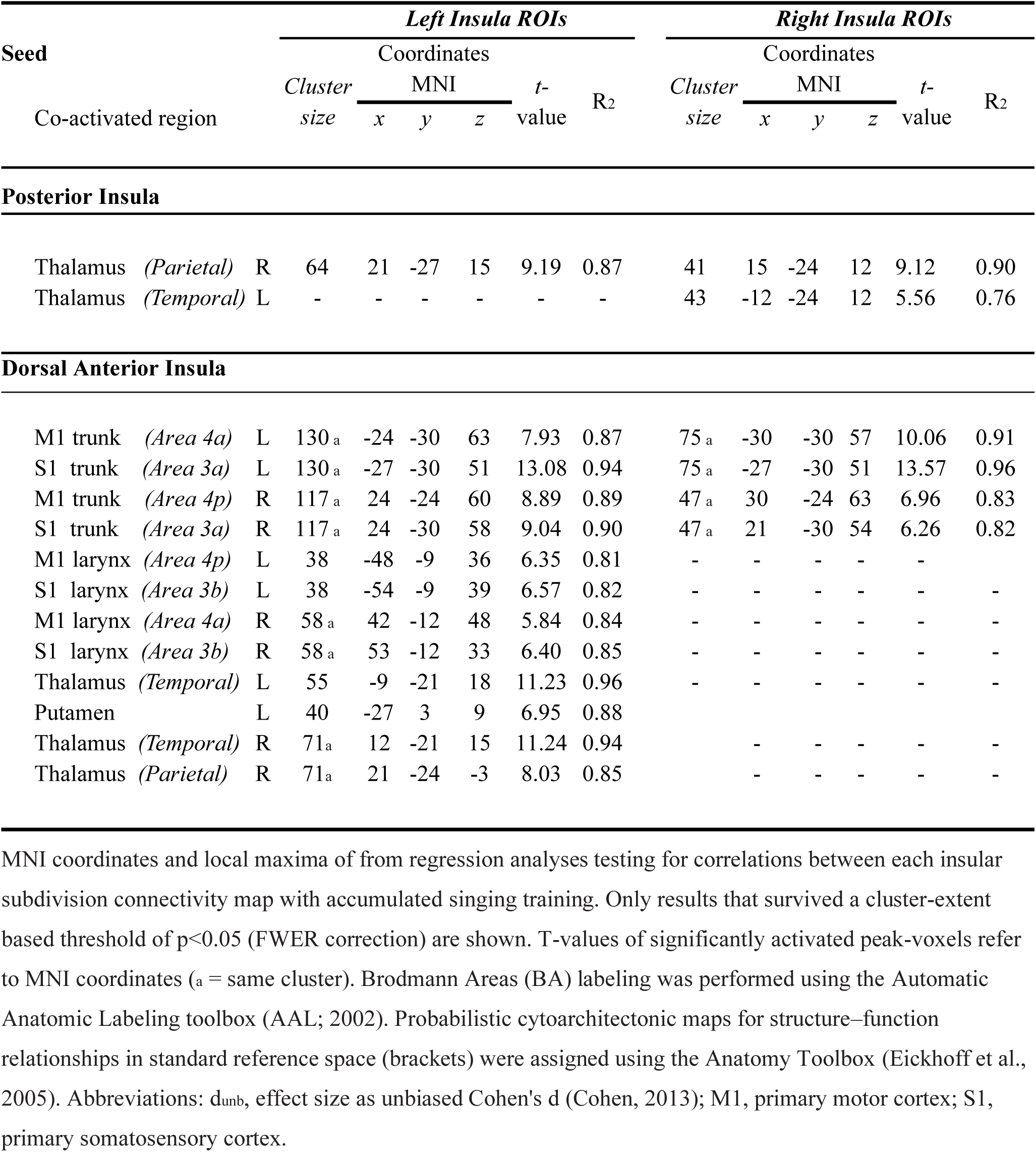
Correlations between insula connectivity maps with accumulated music training

Pitch-matching accuracy revealed no significant correlations with insula connectivity maps across participants. However, accumulated musical training in trained singers was significantly correlated with insula connectivity maps (Figure 3, Table 2):

**Figure 3:**
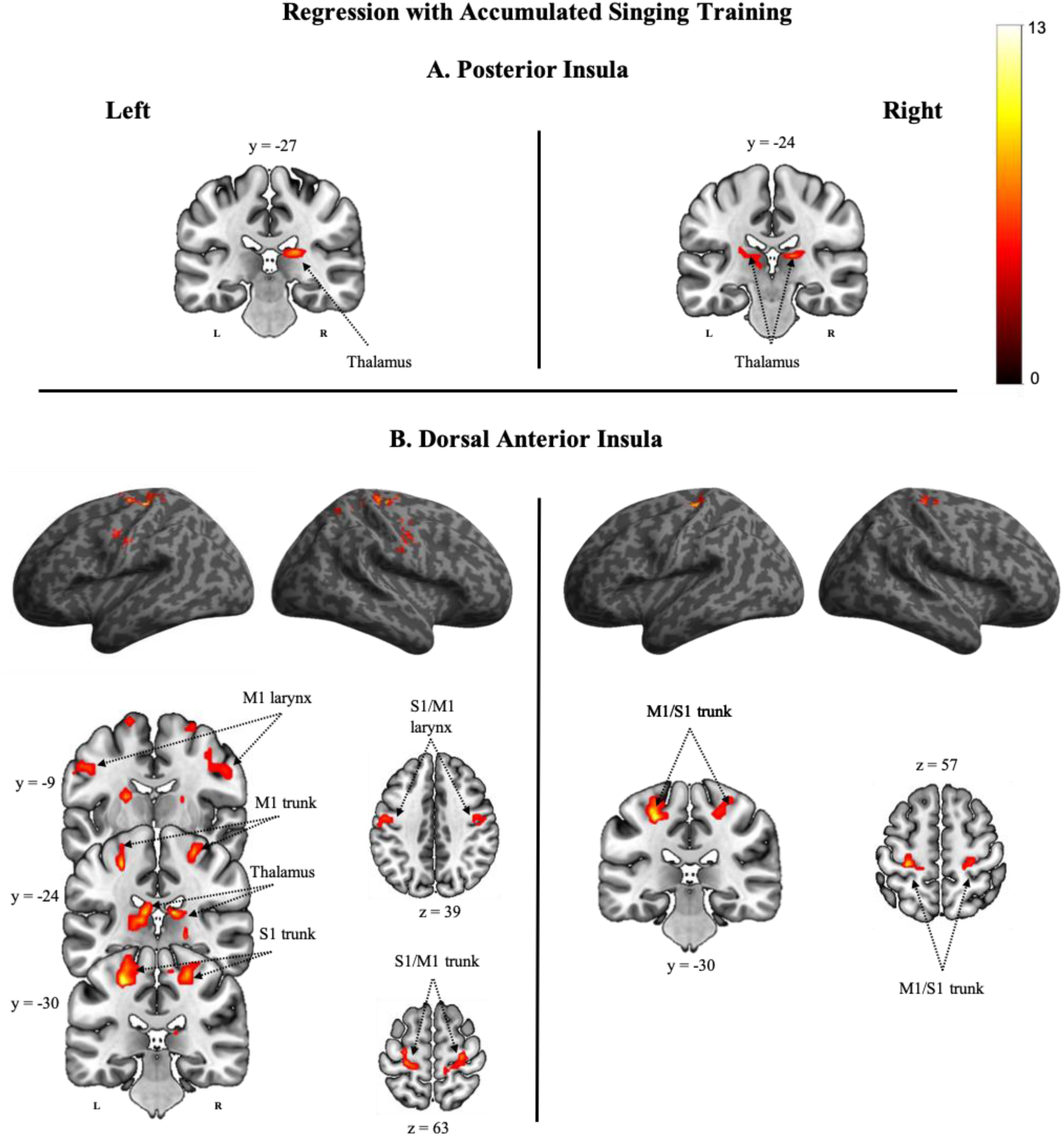
Regression analyses for individual insular ROIs assessing connectivity maps in relation to the amount of accumulated years of vocal training in professional singers. Only results surviving a cluster-extent based significance threshold of p<0.05 (FWER correction) are shown. Detailed information is provided in Table 2. Abbreviations: M1, primary motor cortex; S1, primary somatosensory cortex.

The **left** and **right PI** connectivity maps (Figure 3A) showed a positive correlation with accumulated training in the thalamus.

The **left dAI** connectivity maps (Figure 3B) showed positive correlations with accumulated training in the bilateral representation of the larynx and the diaphragm within M1 and S1, as well as in the bilateral thalamus and the left putamen. The **right dAI** maps showed positive correlations with accumulated training in the bilateral representation of the diaphragm within M1 and S1.

## DISCUSSION

In this resting-state fMRI study, we compared temporally correlated spontaneous blood oxygen level–dependent (BOLD) fluctuations between conservatory trained singers and non-singers. Direct comparisons of insula co-activation patterns revealed increased connectivity patters in singers in bilateral posterior and anterior insula subdivisions with constituents of the sensorimotor speech network. Additionally, accumulated singing training correlated positively with increased connectivity between the dorsal anterior insula with primary motor and somatosensory cortices in the somatotopic representations of the larynx and the diaphragm, the thalamus, and the putamen. These results indicate lasting effects of long-term singing training on insula-based networks involved in speech motor control, suggesting that the integration of intero- and exteroceptive signals of salience that guide fine laryngeal, articulatory, and respiratory sensorimotor control is enhanced in the expert singer.

### Insula connectivity patterns across groups

Previous studies identified a vast number of functional and structural connections between the insula with both cortical and subcortical brain regions subserving sensory, emotional, motivational and cognitive processes (Gogolla, 2017). More specifically, functional segregation of resting-state co-activation patterns identified a tripartite organization of insula subdivisions containing both unique and overlapping functional profiles for the posterior insula (PI), the dorsal anterior insula (dAI), and the ventral anterior insula (vAI) (Uddin et al., 2014), which has been supported by their structural organization (Nomi et al., 2018). The current study confirms the previously reported connectivity patterns (Deen et al., 2011; Zamorano et al., 2017; Zamorano et al., 2019), conforming to a predominantly “cognitive” frontoparietal connectivity pattern for the dAI, a more “affective” co-activation pattern for the vAI, and a more sensorimotor centered co-activation pattern for the PI.

### Group differences in insula connectivity patterns

As predicted, group comparisons revealed enhanced connectivity in trained singers relative to non-singers across all insula subdivisions, whereas the reversed comparison showed no differences.

#### PI connectivity in singers versus non-singers

Conservatory trained singers showed enhanced connectivity between bilateral PI and lobule VI of the cerebellum, a somatotopically organized sensorimotor region that receives somatosensory afferents from the vocal tract and sends efferents to motor areas (Grodd et al., 2001; O’Reilly et al., 2010; Stoodley and Schmahmann, 2010), including those underlying the control of speech and song production (Brown et al., 2006). This is consistent with the unique profile of the PI, which integrates sensorimotor, visceral, and sensory information from the body (Berret et al., 2019; Craig, 2002; Olausson et al., 2002; Pugnaghi et al., 2011) and the vocal tract during speech and orofacial activity (Tsutsumi et al., 2018; Woolnough et al., 2019).

Lesions in PI as well as in the cerebellar lobule VI give rise to speech motor disorders (Ackermann et al., 2007; Baier et al., 2011). Lobule VI shows moreover functional activation during the production and the perception of song and speech (Ackermann, 2008; Callan et al., 2007; Mathiak et al., 2002), thus linking the cerebellum to predictive mechanisms underlying sensorimotor and cognitive aspects of language processing (Argyropoulos, 2016; Friston and Herreros, 2016; Guenther et al., 2006). In this study, increased PI-cerebellar connectivity may therefore reflect enhanced perceptual and prediction processes (Baumann et al., 2015; Moberget and Ivry, 2016) as well as smoother sensorimotor control (Ackermann, 2008), which support enhanced vocal performance in singers (Baer et al., 2015; Kleber et al., 2017).

#### dAI connectivity in singers versus non-singers

Singers also showed enhanced functional connectivity between the left dAI with bilateral inferior parietal lobe (IPL), adjacent S1 (area 2), and the cerebellum (lobule VI). Likewise, the right dAI showed enhanced functional connectivity with right S1 (area 2) and the cerebellum (lobule VI and crus 2). Studies have linked the cerebellum (lobule VI) also to cognitive aspects of language processes (Stoodley et al., 2012), extero- and interoceptive salience processing, and executive control (Crus 2; Baumann et al., 2015; Habas et al., 2009), supported by resting-state data reporting enhanced connectivity among components of the salience network in orchestra musicians (Luo et al., 2014; Zamorano et al., 2017; Zamorano et al., 2019).

The unique functional profile of the dAI suggests indeed a prominent role in salience detection and cognitive processes, including task-related attention switching, inhibition, and error awareness (Menon and Uddin, 2010; Uddin, 2015). Moreover, dAI co-activation patterns comprises brain regions that pertain to phonological and speech processes (Uddin et al., 2014), which links the AI to higher-order aspects of speech motor control (Ackermann and Riecker, 2004, 2010). The latter also requires processing of proprioceptive and body-related information in IPL and S1 (Prevosto et al., 2011) for the preparation and coordination of speech-related vocal tract movements (Bouchard et al., 2013; Nasir and Ostry, 2006; Tremblay et al., 2003). In fact, IPL and S1 contribute nearly seven times more to the vocal motor system in humans than in non-human primates (Kumar et al., 2016). This is reflected in experience-dependent changes in singers, who display enhanced activation in ventral S1, IPL, and the cerebellum during vocal performance compared to non-singers (Kleber et al., 2010), as well as increased gray-matter volume (S1/IPL) (Kleber et al., 2016). An causal involvement of S1 in voice pitch control has been demonstrated with repetitive TMS (Finkel et al., 2019), whereas the dynamic interactions between dAI and S1/IPL in singers facilitate somatosensory feedback integration to compensate for experimentally altered acoustic feedback during singing (Kleber et al., 2017).

Taken together, the enhanced dAI coactivation pattern in singers during rest suggest a fundamental involvement in both sensorimotor and cognitive control pertaining to the execution of vocalizations (Bohland and Guenther, 2006; Eickhoff et al., 2009; Stoodley et al., 2012), perhaps by switching between cognitive and motor aspects of sound-to-speech transformations (Cloutman et al., 2012; Eickhoff et al., 2009; Hickok, 2017; Oh et al., 2014).

#### vAI connectivity in singers versus non-singers

The vAI is generally more strongly associated with valence attribution and affective processing (Craig, 2009, 2010). In the current study, enhanced left vAI co-activation in trained singers encompassed the superior parietal lobe (SPL), intraprietal sulcus (IPS), and the right prefrontal lobe (BA10/frontal pole). The frontal pole mediates limbic reward and learning processes (Denny et al., 2014). The right frontal pole in particular may also support high-level cognitive processes, including language and working memory (Gilbert et al., 2006; Ray et al., 2015). It may therefore represent a gateway to shift attention from sensory monitoring to cue-directed behaviors (Howe et al., 2013), which are crucial features of musical performances.

In contrast, SPL co-activation does not comfortably fit within the vAI’s unique profile but may play role within a frontoparietal attentional control system (Uddin, 2015) that contributes to task performance when stimulus-response associations have been well trained and can be prepared in advance (Corbetta and Shulman, 2002). Increased SPL activation in musicians has been associated with heightened sensorimotor control, increased attention, and working memory load (Foster and Zatorre, 2010; Pallesen et al., 2010). The current findings in singers may therefore reflect enhanced attentional processes related to task-specific awareness (Nelson et al., 2010).

### Insula connectivity as a function of accumulated singing training

In terms of functional anatomy, the insula contributes to the spinothalamocortical pathway, in which the lamina I projects afferent information from body tissues via the thalamus to both the somatosensory area 3a and the posterior insula, the so-called interoceptive cortex (Craig, 2002, 2009, 2010). These signals are then re-represented, integrated, and affectively evaluated along a posterior-to-anterior insula progression scheme, thereby generating a comprehensive representation of one’s emotional responses. The enhanced connectivity patterns found as a function of singing experience in this study encompassed the constituents of this pathway. Specifically, between (i) the bilateral PI with the thalamus, (ii) the bilateral dAI with the sensorimotor representation (S1/M1) of the diaphragm, and (iii) the left dAI with the bilateral sensorimotor representation (S1/M1) of the larynx, thalamus (bilaterally), and left putamen. The precise functional association of these co-activated regions in singers (i) confirms the insula as a major functional integration site for task-relevant bodily information and (ii) suggests that singing training may enhance access to these resources (Kleber et al., 2017; Schirmer-Mokwa et al., 2015; Zamorano et al., 2017; Zamorano et al., 2019).

That is, the coactivated regions process visceral and somatosensory information underlying respiratory, laryngeal, and articulatory actions (Banzett et al., 2000; Bouchard et al., 2013; Conant et al., 2018; Pfenning et al., 2014) underlying vocalization related movements, which are accompanied by proprioceptive, kinesthetic, and interoceptive signals that are integrated with the motor system to guide song and speech production (Simonyan and Horwitz, 2011; Smotherman, 2007; Tremblay et al., 2003). More specifically, individual movements stimulate a corresponding somatosensory region in S1 (Giraud and Poeppel, 2012), which is intrinsically connected to ipsilateral M1 and IPL (Bouchard et al., 2013) and becomes tightly associated with auditory-motor transformations through experience (Ito et al., 2016). M1 sends its strongest subcortical projections to the putamen (Simonyan and Horwitz, 2011), which shows enhanced task-specific activation in proficient singers (Kleber et al., 2010; Segado et al., 2018).

The insula is known to connect with cortical areas involved in the sensorimotor and cognitive control of speech (Battistella et al., 2018), supposedly by integrating sensory and visceral inputs that subserve the coordination of both respiratory and vocal tract musculature (Ackermann and Riecker, 2010; Dronkers, 1996; Oh et al., 2014). Interestingly, the cortical areas that control the interaction of expiratory, laryngeal, and articulatory muscle groups during planned vocalizations are somatotopically overlapping (Belyk and Brown, 2017; Kumar et al., 2016; Loucks et al., 2007). For example, voluntary inspiration (e.g., taking a deeper than normal breath before a long musical phrase) is more dorsally located within the somatotopic representation of the trunk and diaphragm (Eickhoff et al., 2008), whereas expiration in the context of vocalization is in comparison more ventrally located (Belyk and Brown, 2017). As both of these regions are more active in trained singers than in non-singers during singing (Kleber et al., 2017; Kleber et al., 2013), increased insular connectivity with these areas may support vocal performance through enhanced integration of vocalization-related bodily signals (Kleber et al., 2017; Kleber et al., 2013; Schirmer-Mokwa et al., 2015; Zarate, 2013).

### Limitations

One limitation of this study includes the fact that fMRI exposes participants to continuous scanner noise, which can reduce the robustness and the replicability of functional connectivity findings within the somatosensory, auditory, and motor networks (Andoh et al., 2017). However, robust experience-dependent differences in S1/M1 in vocalization areas in the current study replicate previous findings with classical instrumental musicians, in which greater experience yielded greater insular connectivity with the sensorimotor hand area (Zamorano et al., 2017). This specificity with regards to functional correspondence increases the confidence in the current results and supports the notion that insula-based networks critically support body awareness underlying music production. Regarding the generalization of our results, we must consider the moderate sample size and that all singers were conservatory trained. Therefore, we cannot rule out the possibility that co-activation patterns may differ in amateur-level singers and suggest that future studies should replicate these results in larger samples.

### Conclusions

This study examined the impact of conservatory-level singing training on insula-based network connectivity at rest. The results of this research support a role of the insula in modulating sensory integration within vocalization related respiratory and laryngeal systems (Ackermann and Riecker, 2010; Eickhoff et al., 2009), perhaps by regulating conscious and non-conscious aspects of salience processing associated with task-relevant bodily processes (Kleber et al., 2017). Moreover, we propose that the insula’s functional involvement in the processing, manipulation, and transformation of sensory information can be altered through experience, based on the notion that neuroplastic changes associated with multisensory integration during musical training involve the insula in task general and task-specific manners (Luo et al., 2012; Palomar-Garcia et al., 2017; Zamorano et al., 2017; Zamorano et al., 2019). Together, these data add to a growing body of literature suggesting that trained singers rely increasingly on bodily feedback processes to support vocal motor control (Jones and Keough, 2008; Kleber et al., 2017; Kleber et al., 2010; Kleber et al., 2013; Mürbe et al., 2004; Zarate and Zatorre, 2008), in line with reports on heightened sensitivity to interoceptive and exteroceptive stimuli in performing artists (Christensen et al., 2017; Schirmer-Mokwa et al., 2015; Zamorano et al., 2014).

---

**S-Figure 1:**
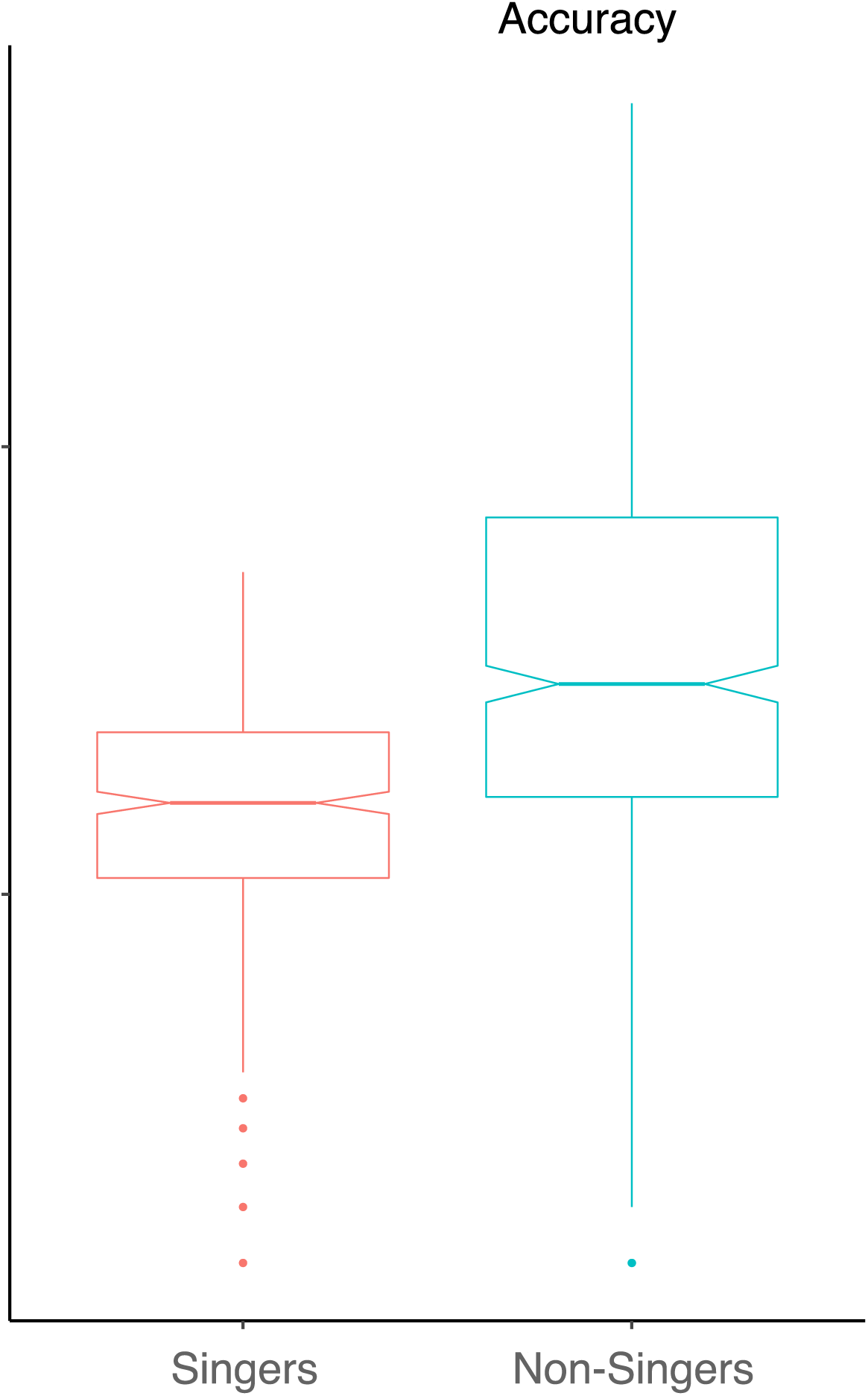
Pitch-matching accuracy in singers and non-singers, representing the deviation between the target tones and the tones sung by the participants. Deviation was measured in cents (one semitone corresponds to 100 cents). As the residuals were found to be non-normal and there was evidence of heteroscedasticity of variances, pitch-deviation data were log transformed (log10) prior to statistical analysis.

## Acknowledgments

We would like to thank Sebastian Finkel for assistance in data acquisition and Ben Deen for providing the insula subdivision ROIs from his earlier work (https://bendeen.com/data/). This study was supported by a grant from the Deutsche Forschungsgemeinschaft to B.K. (KL 2341/1-1). The Center for Music in the Brain is funded by the Danish National Research Foundation (DNRF 117).

## References

Ackermann, H., 2008. Cerebellar contributions to speech production and speech perception: psycholinguistic and neurobiological perspectives. Trends Neurosci 31, 265–272.

Ackermann, H., Mathiak, K., Riecker, A., 2007. The contribution of the cerebellum to speech production and speech perception: clinical and functional imaging data. Cerebellum (London, England) 6, 202–213.

Ackermann, H., Riecker, A., 2004. The contribution of the insula to motor aspects of speech production: a review and a hypothesis. Brain Lang 89, 320–328.

Ackermann, H., Riecker, A., 2010. The contribution(s) of the insula to speech production: a review of the clinical and functional imaging literature. Brain Struct Funct 214, 419–433.

Albouy, P., Benjamin, L., Morillon, B., Zatorre, R.J., 2020. Distinct sensitivity to spectrotemporal modulation supports brain asymmetry for speech and melody. Science 367, 1043–1047.

Andoh, J., Ferreira, M., Leppert, I.R., Matsushita, R., Pike, B., Zatorre, R.J., 2017. How restful is it with all that noise? Comparison of Interleaved silent steady state (ISSS) and conventional imaging in resting-state fMRI. NeuroImage 147, 726–735.

Ardila, A., Bernal, B., Rosselli, M., 2014. Participation of the insula in language revisited: A meta-analytic connectivity study. J Neurolinguist 29, 31–41.

Argyropoulos, G.P., 2016. The cerebellum, internal models and prediction in ‘non-motor’ aspects of language: A critical review. Brain Lang 161, 4–17.

Ashburner, J., Friston, K.J., 2005. Unified segmentation. NeuroImage 26, 839–851.

Baer, L.H., Park, M.T., Bailey, J.A., Chakravarty, M.M., Li, K.Z., Penhune, V.B., 2015. Regional cerebellar volumes are related to early musical training and finger tapping performance. NeuroImage 109, 130–139.

Baier, B., zu Eulenburg, P., Glassl, O., Dieterich, M., 2011. Lesions to the posterior insular cortex cause dysarthria. European journal of neurology : the official journal of the European Federation of Neurological Societies 18, 1429–1431.

Banzett, R.B., Mulnier, H.E., Murphy, K., Rosen, S.D., Wise, R.J., Adams, L., 2000. Breathlessness in humans activates insular cortex. NeuroReport 11, 2117–2120.

Barr, D.J., Levy, R., Scheepers, C., Tily, H.J., 2013. Random effects structure for confirmatory hypothesis testing: Keep it maximal. J Mem Lang 68.

Bates, D., Mächler, M., Bolker, B., Walker, S., 2015. Fitting Linear Mixed-Effects Models Using lme4. Journal of Statistical Software 67.

Battistella, G., Kumar, V., Simonyan, K., 2018. Connectivity profiles of the insular network for speech control in healthy individuals and patients with spasmodic dysphonia. Brain Struct Funct 223, 2489–2498.

Baumann, O., Borra, R.J., Bower, J.M., Cullen, K.E., Habas, C., Ivry, R.B., Leggio, M., Mattingley, J.B., Molinari, M., Moulton, E.A., Paulin, M.G., Pavlova, M.A., Schmahmann, J.D., Sokolov, A.A., 2015. Consensus paper: the role of the cerebellum in perceptual processes. Cerebellum (London, England) 14, 197–220.

Belyk, M., Brown, S., 2017. The origins of the vocal brain in humans. Neurosci Biobehav Rev 77, 177–193.

Berret, E., Kintscher, M., Palchaudhuri, S., Tang, W., Osypenko, D., Kochubey, O., Schneggenburger, R., 2019. Insular cortex processes aversive somatosensory information and is crucial for threat learning. Science 364.

Bohland, J.W., Guenther, F.H., 2006. An fMRI investigation of syllable sequence production. NeuroImage 32, 821–841.

Bouchard, K.E., Mesgarani, N., Johnson, K., Chang, E.F., 2013. Functional organization of human sensorimotor cortex for speech articulation. Nature 495, 327–332.

Brown, S., Martinez, M.J., Parsons, L.M., 2006. Music and language side by side in the brain: a PET study of the generation of melodies and sentences. Eur J Neurosci 23, 2791–2803.

Callan, D.E., Kawato, M., Parsons, L., Turner, R., 2007. Speech and song: the role of the cerebellum. Cerebellum (London, England) 6, 321–327.

Chao-Gan, Y., Yu-Feng, Z., 2010. DPARSF: A MATLAB Toolbox for “Pipeline” Data Analysis of Resting-State fMRI. Frontiers in systems neuroscience 4, 13.

Christensen, J.F., Gaigg, S.B., Calvo-Merino, B., 2017. I can feel my heartbeat: Dancers have increased interoceptive accuracy. Psychophysiology.

Cloutman, L.L., Binney, R.J., Drakesmith, M., Parker, G.J., Lambon Ralph, M.A., 2012. The variation of function across the human insula mirrors its patterns of structural connectivity: evidence from in vivo probabilistic tractography. NeuroImage 59, 3514–3521.

Cohen, J., 2013. Statistical power analysis for the behavioral sciences. Academic press.

Conant, D.F., Bouchard, K.E., Leonard, M.K., Chang, E.F., 2018. Human Sensorimotor Cortex Control of Directly Measured Vocal Tract Movements during Vowel Production. J Neurosci 38, 2955–2966.

Corbetta, M., Shulman, G.L., 2002. Control of goal-directed and stimulus-driven attention in the brain. Nat Rev Neurosci 3, 201–215.

Cox, R.W., 1996. AFNI: software for analysis and visualization of functional magnetic resonance neuroimages. Computers and biomedical research, an international journal 29, 162–173.

Craig, A.D., 2002. How do you feel? Interoception: the sense of the physiological condition of the body. Nat Rev Neurosci 3, 655–666.

Craig, A.D., 2009. How do you feel--now? The anterior insula and human awareness. Nat Rev Neurosci 10, 59–70.

Craig, A.D., 2010. The sentient self. Brain Struct Funct 214, 563–577.

Deen, B., Pitskel, N.B., Pelphrey, K.A., 2011. Three systems of insular functional connectivity identified with cluster analysis. Cereb Cortex 21, 1498–1506.

Denny, B.T., Ochsner, K.N., Weber, J., Wager, T.D., 2014. Anticipatory brain activity predicts the success or failure of subsequent emotion regulation. Social cognitive and affective neuroscience 9, 403–411.

Dick, A.S., Bernal, B., Tremblay, P., 2014. The language connectome: new pathways, new concepts. Neuroscientist 20, 453–467.

Donahue, E.N., Leborgne, W.D., Brehm, S.B., Weinrich, B.D., 2014. Reported vocal habits of first-year undergraduate musical theater majors in a preprofessional training program: a 10-year retrospective study. J Voice 28, 316–323.

Dronkers, N.F., 1996. A new brain region for coordinating speech articulation. Nature 384, 159–161.

Eickhoff, S.B., Grefkes, C., Fink, G.R., Zilles, K., 2008. Functional lateralization of face, hand, and trunk representation in anatomically defined human somatosensory areas. Cereb Cortex 18, 2820–2830.

Eickhoff, S.B., Heim, S., Zilles, K., Amunts, K., 2009. A systems perspective on the effective connectivity of overt speech production. Philosophical transactions. Series A, Mathematical, physical, and engineering sciences 367, 2399–2421.

Eickhoff, S.B., Stephan, K.E., Mohlberg, H., Grefkes, C., Fink, G.R., Amunts, K., Zilles, K., 2005. A new SPM toolbox for combining probabilistic cytoarchitectonic maps and functional imaging data. NeuroImage 25, 1325–1335.

Finkel, S., Veit, R., Lotze, M., Friberg, A., Vuust, P., Soekadar, S., Birbaumer, N., Kleber, B., 2019. Intermittent theta burst stimulation over right somatosensory larynx cortex enhances vocal pitch-regulation in nonsingers. Hum Brain Mapp 40, 2174–2187.

Foster, N.E., Zatorre, R.J., 2010. A role for the intraparietal sulcus in transforming musical pitch information. Cereb Cortex 20, 1350–1359.

Fox, M.D., Raichle, M.E., 2007. Spontaneous fluctuations in brain activity observed with functional magnetic resonance imaging. Nat Rev Neurosci 8, 700–711.

Friberg, A., Schoonderwaldt, E., Juslin, P.N., 2005. CUEX: An algorithm for extracting expressive tone variables from audio recordings. Acoustica united with Acta Acoustica 93, 411–420.

Friston, K., Herreros, I., 2016. Active Inference and Learning in the Cerebellum. Neural Comput 28, 1812–1839.

Gilbert, S.J., Spengler, S., Simons, J.S., Steele, J.D., Lawrie, S.M., Frith, C.D., Burgess, P.W., 2006. Functional specialization within rostral prefrontal cortex (area 10): a meta-analysis. J Cogn Neurosci 18, 932–948.

Giraud, A.L., Poeppel, D., 2012. Cortical oscillations and speech processing: emerging computational principles and operations. Nat Neurosci 15, 511–517.

Gogolla, N., 2017. The insular cortex. Curr Biol 27, R580–R586.

Grodd, W., Hulsmann, E., Lotze, M., Wildgruber, D., Erb, M., 2001. Sensorimotor mapping of the human cerebellum: fMRI evidence of somatotopic organization. Hum Brain Mapp 13, 55–73.

Guenther, F.H., 2016. Neural control of speech. The MIT Press, Cambridge, MA.

Guenther, F.H., Ghosh, S.S., Tourville, J.A., 2006. Neural modeling and imaging of the cortical interactions underlying syllable production. Brain Lang 96, 280–301.

Guenther, F.H., Hickok, G., 2015. Role of the auditory system in speech production. Handb Clin Neurol 129, 161–175.

Guerra-Carrillo, B., Mackey, A.P., Bunge, S.A., 2014. Resting-state fMRI: a window into human brain plasticity. Neuroscientist 20, 522–533.

Habas, C., Kamdar, N., Nguyen, D., Prater, K., Beckmann, C.F., Menon, V., Greicius, M.D., 2009. Distinct cerebellar contributions to intrinsic connectivity networks. J Neurosci 29, 8586–8594.

Hage, S.R., Nieder, A., 2016. Dual Neural Network Model for the Evolution of Speech and Language. Trends Neurosci 39, 813–829.

Heller, R., Stanley, D., Yekutieli, D., Rubin, N., Benjamini, Y., 2006. Cluster-based analysis of FMRI data. NeuroImage 33, 599–608.

Herholz, S.C., Zatorre, R.J., 2012. Musical training as a framework for brain plasticity: behavior, function, and structure. Neuron 76, 486–502.

Hertrich, I., Dietrich, S., Ackermann, H., 2016. The role of the supplementary motor area for speech and language processing. Neurosci Biobehav Rev 68, 602–610.

Hickok, G., 2017. A cortical circuit for voluntary laryngeal control: Implications for the evolution language. Psychon Bull Rev 24, 56–63.

Howe, W.M., Berry, A.S., Francois, J., Gilmour, G., Carp, J.M., Tricklebank, M., Lustig, C., Sarter, M., 2013. Prefrontal cholinergic mechanisms instigating shifts from monitoring for cues to cue-guided performance: converging electrochemical and fMRI evidence from rats and humans. J Neurosci 33, 8742–8752.

Ito, T., Coppola, J.H., Ostry, D.J., 2016. Speech motor learning changes the neural response to both auditory and somatosensory signals. Sci Rep 6, 25926.

Jeffries, K.J., Fritz, J.B., Braun, A.R., 2003. Words in melody: an H(2)15O PET study of brain activation during singing and speaking. NeuroReport 14, 749–754.

Jones, J.A., Keough, D., 2008. Auditory-motor mapping for pitch control in singers and nonsingers. Exp Brain Res 190, 279–287.

Jurgens, U., 2002. Neural pathways underlying vocal control. Neurosci Biobehav Rev 26, 235–258.

Jurgens, U., 2009. The neural control of vocalization in mammals: a review. J Voice 23, 1–10.

Kleber, B., Friberg, A., Zeitouni, A., Zatorre, R., 2017. Experience-dependent modulation of right anterior insula and sensorimotor regions as a function of noise-masked auditory feedback in singers and nonsingers. NeuroImage 147, 97–110.

Kleber, B., Veit, R., Birbaumer, N., Gruzelier, J., Lotze, M., 2010. The brain of opera singers: experience-dependent changes in functional activation. Cereb Cortex 20, 1144–1152.

Kleber, B., Veit, R., Moll, C.V., Gaser, C., Birbaumer, N., Lotze, M., 2016. Voxel-based morphometry in opera singers: Increased gray-matter volume in right somatosensory and auditory cortices. NeuroImage 133, 477–483.

Kleber, B., Zarate, J.M., 2014. The Neuroscience of Singing. In: Graham, W., Nix, J. (Eds.), The Oxford Handbook of Singing. Oxford University Press, Oxford, UK.

Kleber, B., Zeitouni, A.G., Friberg, A., Zatorre, R.J., 2013. Experience-dependent modulation of feedback integration during singing: role of the right anterior insula. J Neurosci 33, 6070–6080.

Klein, C., Liem, F., Hanggi, J., Elmer, S., Jancke, L., 2016. The “silent” imprint of musical training. Hum Brain Mapp 37, 536–546.

Kumar, V., Croxson, P.L., Simonyan, K., 2016. Structural Organization of the Laryngeal Motor Cortical Network and Its Implication for Evolution of Speech Production. J Neurosci 36, 4170–4181.

Loucks, T.M., Poletto, C.J., Simonyan, K., Reynolds, C.L., Ludlow, C.L., 2007. Human brain activation during phonation and exhalation: common volitional control for two upper airway functions. NeuroImage 36, 131–143.

Luo, C., Guo, Z.W., Lai, Y.X., Liao, W., Liu, Q., Kendrick, K.M., Yao, D.Z., Li, H., 2012. Musical training induces functional plasticity in perceptual and motor networks: insights from resting-state FMRI. PLoS ONE 7, e36568.

Luo, C., Tu, S., Peng, Y., Gao, S., Li, J., Dong, L., Li, G., Lai, Y., Li, H., Yao, D., 2014. Long-term effects of musical training and functional plasticity in salience system. Neural Plast 2014, 180138.

Mathiak, K., Hertrich, I., Grodd, W., Ackermann, H., 2002. Cerebellum and speech perception: a functional magnetic resonance imaging study. J Cogn Neurosci 14, 902–912.

Menon, V., Uddin, L.Q., 2010. Saliency, switching, attention and control: a network model of insula function. Brain Struct Funct 214, 655–667.

Moberget, T., Ivry, R.B., 2016. Cerebellar contributions to motor control and language comprehension: searching for common computational principles. Ann N Y Acad Sci 1369, 154–171.

Munte, T.F., Altenmüller, E., Jancke, L., 2002. The musician’s brain as a model of neuroplasticity. Nat Rev Neurosci 3, 473–478.

Mürbe, D., Pabst, F., Hofmann, G., Sundberg, J., 2004. Effects of a professional solo singer education on auditory and kinesthetic feedback--a longitudinal study of singers’ pitch control. Journal of Voice 18, 236–241.

Murphy, K., Birn, R.M., Handwerker, D.A., Jones, T.B., Bandettini, P.A., 2009. The impact of global signal regression on resting state correlations: are anti-correlated networks introduced? NeuroImage 44, 893–905.

Nasir, S.M., Ostry, D.J., 2006. Somatosensory precision in speech production. Curr Biol 16, 1918–1923.

Nelson, S.M., Dosenbach, N.U., Cohen, A.L., Wheeler, M.E., Schlaggar, B.L., Petersen, S.E., 2010. Role of the anterior insula in task-level control and focal attention. Brain Struct Funct 214, 669–680.

Nomi, J.S., Schettini, E., Broce, I., Dick, A.S., Uddin, L.Q., 2018. Structural Connections of Functionally Defined Human Insular Subdivisions. Cereb Cortex 28, 3445–3456.

O’Reilly, J.X., Beckmann, C.F., Tomassini, V., Ramnani, N., Johansen-Berg, H., 2010. Distinct and overlapping functional zones in the cerebellum defined by resting state functional connectivity. Cereb Cortex 20, 953–965.

Oh, A., Duerden, E.G., Pang, E.W., 2014. The role of the insula in speech and language processing. Brain Lang 135, 96–103.

Olausson, H., Lamarre, Y., Backlund, H., Morin, C., Wallin, B.G., Starck, G., Ekholm, S., Strigo, I., Worsley, K., Vallbo, A.B., Bushnell, M.C., 2002. Unmyelinated tactile afferents signal touch and project to insular cortex. Nat Neurosci 5, 900–904.

Pallesen, K.J., Brattico, E., Bailey, C.J., Korvenoja, A., Koivisto, J., Gjedde, A., Carlson, S., 2010. Cognitive control in auditory working memory is enhanced in musicians. PLoS ONE 5, e11120.

Palomar-Garcia, M.A., Zatorre, R.J., Ventura-Campos, N., Bueicheku, E., Avila, C., 2017. Modulation of Functional Connectivity in Auditory-Motor Networks in Musicians Compared with Nonmusicians. Cereb Cortex 27, 2768–2778.

Parrell, B., Lammert, A.C., Ciccarelli, G., Quatieri, T.F., 2019. Current models of speech motor control: A control-theoretic overview of architectures and properties. J Acoust Soc Am 145, 1456.

Pfenning, A.R., Hara, E., Whitney, O., Rivas, M.V., Wang, R., Roulhac, P.L., Howard, J.T., Wirthlin, M., Lovell, P.V., Ganapathy, G., Mouncastle, J., Moseley, M.A., Thompson, J.W., Soderblom, E.J., Iriki, A., Kato, M., Gilbert, M.T., Zhang, G., Bakken, T., Bongaarts, A., Bernard, A., Lein, E., Mello, C.V., Hartemink, A.J., Jarvis, E.D., 2014. Convergent transcriptional specializations in the brains of humans and song-learning birds. Science 346, 1256846.

Prevosto, V., Graf, W., Ugolini, G., 2011. Proprioceptive pathways to posterior parietal areas MIP and LIPv from the dorsal column nuclei and the postcentral somatosensory cortex. Eur J Neurosci 33, 444–460.

Price, C.J., 2000. The anatomy of language: contributions from functional neuroimaging. Journal of anatomy 197 Pt 3, 335–359.

Price, C.J., 2012. A review and synthesis of the first 20 years of PET and fMRI studies of heard speech, spoken language and reading. NeuroImage 62, 816–847.

Pugnaghi, M., Meletti, S., Castana, L., Francione, S., Nobili, L., Mai, R., Tassi, L., 2011. Features of somatosensory manifestations induced by intracranial electrical stimulations of the human insula. Clin Neurophysiol 122, 2049–2058.

Ray, K.L., Zald, D.H., Bludau, S., Riedel, M.C., Bzdok, D., Yanes, J., Falcone, K.E., Amunts, K., Fox, P.T., Eickhoff, S.B., Laird, A.R., 2015. Co-activation based parcellation of the human frontal pole. NeuroImage 123, 200–211.

Remedios, R., Logothetis, N.K., Kayser, C., 2009. An auditory region in the primate insular cortex responding preferentially to vocal communication sounds. J Neurosci 29, 1034–1045.

Riecker, A., Ackermann, H., Wildgruber, D., Dogil, G., Grodd, W., 2000. Opposite hemispheric lateralization effects during speaking and singing at motor cortex, insula and cerebellum. NeuroReport 11, 1997–2000.

Riecker, A., Brendel, B., Ziegler, W., Erb, M., Ackermann, H., 2008. The influence of syllable onset complexity and syllable frequency on speech motor control. Brain Lang 107, 102–113.

Schirmer-Mokwa, K.L., Fard, P.R., Zamorano, A.M., Finkel, S., Birbaumer, N., Kleber, B.A., 2015. Evidence for Enhanced Interoceptive Accuracy in Professional Musicians. Front Behav Neurosci 9, 349.

Schlaug, G., 2015. Musicians and music making as a model for the study of brain plasticity. Progress in brain research 217, 37–55.

Seeley, W.W., Menon, V., Schatzberg, A.F., Keller, J., Glover, G.H., Kenna, H., Reiss, A.L., Greicius, M.D., 2007. Dissociable intrinsic connectivity networks for salience processing and executive control. J Neurosci 27, 2349–2356.

Segado, M., Hollinger, A., Thibodeau, J., Penhune, V., Zatorre, R.J., 2018. Partially Overlapping Brain Networks for Singing and Cello Playing. Front Neurosci 12, 351.

Simonyan, K., Horwitz, B., 2011. Laryngeal motor cortex and control of speech in humans. Neuroscientist 17, 197–208.

Singer, T., Critchley, H.D., Preuschoff, K., 2009. A common role of insula in feelings, empathy and uncertainty. Trends Cogn Sci 13, 334–340.

Smith, A., 2006. Speech motor development: Integrating muscles, movements, and linguistic units. J Commun Disord 39, 331–349.

Smotherman, M.S., 2007. Sensory feedback control of mammalian vocalizations. Behav Brain Res 182, 315–326.

Stadler Elmer, S., 2011. Human singing: Towards a developmental theory. Psychomusicology 21, 13–30.

Stoodley, C.J., Schmahmann, J.D., 2010. Evidence for topographic organization in the cerebellum of motor control versus cognitive and affective processing. Cortex 46, 831–844.

Stoodley, C.J., Valera, E.M., Schmahmann, J.D., 2012. Functional topography of the cerebellum for motor and cognitive tasks: an fMRI study. NeuroImage 59, 1560–1570.

Strigo, I.A., Craig, A.D., 2016. Interoception, homeostatic emotions and sympathovagal balance. Philosophical transactions of the Royal Society of London 371.

Tremblay, S., Shiller, D.M., Ostry, D.J., 2003. Somatosensory basis of speech production. Nature 423, 866–869.

Tsutsumi, Y., Tachibana, Y., Sato, F., Furuta, T., Ohara, H., Tomita, A., Fujita, M., Moritani, M., Yoshida, A., 2018. Cortical and Subcortical Projections from Granular Insular Cortex Receiving Orofacial Proprioception. Neuroscience 388, 317–329.

Tzourio-Mazoyer, N., Landeau, B., Papathanassiou, D., Crivello, F., Etard, O., Delcroix, N., Mazoyer, B., Joliot, M., 2002. Automated anatomical labeling of activations in SPM using a macroscopic anatomical parcellation of the MNI MRI single-subject brain. NeuroImage 15, 273–289.

Uddin, L.Q., 2015. Salience processing and insular cortical function and dysfunction. Nat Rev Neurosci 16, 55–61.

Uddin, L.Q., Kinnison, J., Pessoa, L., Anderson, M.L., 2014. Beyond the tripartite cognition-emotion-interoception model of the human insular cortex. J Cogn Neurosci 26, 16–27.

Vuust, P., Gebauer, L.K., Witek, M.A., 2014. Neural underpinnings of music: the polyrhythmic brain. Advances in experimental medicine and biology 829, 339–356.

Weiss-Croft, L.J., Baldeweg, T., 2015. Maturation of language networks in children: A systematic review of 22years of functional MRI. NeuroImage 123, 269–281.

Woo, C.W., Krishnan, A., Wager, T.D., 2014. Cluster-extent based thresholding in fMRI analyses: pitfalls and recommendations. NeuroImage 91, 412–419.

Woolnough, O., Forseth, K.J., Rollo, P.S., Tandon, N., 2019. Uncovering the functional anatomy of the human insula during speech. Elife 8.

Yan, C.G., Wang, X.D., Zuo, X.N., Zang, Y.F., 2016. DPABI: Data Processing & Analysis for (Resting-State) Brain Imaging. Neuroinformatics 14, 339–351.

Zamorano, A.M., Cifre, I., Montoya, P., Riquelme, I., Kleber, B., 2017. Insula-based networks in professional musicians: Evidence for increased functional connectivity during resting state fMRI. Hum Brain Mapp 38, 4834–4849.

Zamorano, A.M., Montoya, P., Cifre, I., Vuust, P., Riquelme, I., Kleber, B., 2019. Experience-dependent neuroplasticity in trained musicians modulates the effects of chronic pain on insula-based networks - A resting-state fMRI study. NeuroImage 202, 116103.

Zamorano, A.M., Riquelme, I., Kleber, B., Altenmüller, E., Hatem, S.M., Montoya, P., 2014. Pain sensitivity and tactile spatial acuity are altered in healthy musicians as in chronic pain patients. Front Hum Neurosci 8, 1016.

Zarate, J.M., 2013. The neural control of singing. Front Hum Neurosci 7, 237.

Zarate, J.M., Zatorre, R.J., 2008. Experience-dependent neural substrates involved in vocal pitch regulation during singing. NeuroImage 40, 1871–1887.

Zatorre, R.J., 2013. Predispositions and plasticity in music and speech learning: neural correlates and implications. Science 342, 585–589.

